# X-intNMF: A Cross- and Intra-Omics Regularized NMF Framework for Multi-Omics Integration

**DOI:** 10.1101/2025.09.17.676796

**Authors:** Tien-Thanh Bui, Rui Xie, Wei Zhang

## Abstract

**Motivation:** The rapid accumulation of multi-omics data presents a valuable opportunity to advance our understanding of complex diseases and biological systems, driving the development of integrative computational methods. However, the complexity of biological processes—spanning multiple molecular layers and involving intricate regulatory interactions—requires models that can capture both intra- and cross-omics relationships. Most existing integration methods primarily focus on sample-level similarities or intra-omics feature interactions, often neglecting the interactions across different omics layers. This limitation can result in the loss of critical biological information and suboptimal performance. To address this gap, we propose X-intNMF, a network-regularized non-negative matrix factorization (NMF) model that explicitly incorporates both cross-omics and intra-omics feature interaction networks during multi-omics integration. By modeling these multi-layered relationships, X-intNMF enhances the representation of biological interactions and improves integration quality and prediction accuracy.

**Results:** For evaluation, we applied X-intNMF to predict breast cancer phenotypes and classify clinical outcomes in lung and ovarian cancers using mRNA expression, microRNA expression, and DNA methylation data from TCGA. The results show that X-intNMF consistently outperforms state-of-the-art methods. Ablation studies confirm that incorporating both cross-omics and intra-omics interactions contributes significantly to the model’s improved performance. Additionally, survival analysis demonstrates that the integrated multi-omics representation offers strong prognostic value for both overall survival and disease-free status. These findings highlight X-intNMF’s ability to effectively model multi-layered molecular interactions while maintaining interpretability, robustness, and scalability within the NMF framework.

**Availability and implementation:** Source code is available at: https://github.com/compbiolabucf/X-intNMF

## Introduction

Advancements in high-throughput sequencing technologies have enabled the large-scale collection of omics data across diverse molecular layers, with each modality capturing a distinct aspect of biological regulation. However, the inherently multi-layered and complex nature of biological systems means that analyzing any single omics type in isolation often leads to an incomplete understanding. To gain a more comprehensive and accurate view of biological processes, it is essential to integrate data from multiple omics layers. As a result, multi-omics integration has emerged as a critical area in biomedical research, aiming to combine heterogeneous molecular data to better characterize biological systems [1]. This integrative strategy not only deepens our insight into biological complexity but also enables a wide range of biomedical applications, such as precision oncology, personalized medicine, and microbiome research [2, 3].

Despite the advantages of multi-omics integration, several key challenges remain, including the high dimensionality within individual omics layers, heterogeneity across data distributions, limited interpretability, and scalability issues when processing multi-modal data. To address these challenges, a wide range of computational approaches have been developed, including kernel-based methods, probabilistic models [4], matrix factorization techniques [5, 6], and deep learning frameworks [7, 8, 9]. Matrix factorization and probabilistic approaches are commonly used for their simplicity and interpretability, but they often suffer from high computational costs and limited scalability. In contrast, deep learning methods have demonstrated strong performance in multi-omics integration tasks but typically require large training datasets and often lack transparency and interpretability. Furthermore, many existing methods are designed for specific omics types, limiting their generalizability and flexibility when applied to heterogeneous datasets. Notably, biological interaction networks, particularly cross-omics interactions such as mRNA–miRNA regulatory relationships, are frequently neglected. Overlooking these interactions can lead to substantial information loss and diminished model performance, despite their critical role in understanding complex regulatory mechanisms [10].

Matrix factorization, which decomposes an input matrix into the product of two or more low-rank matrices, is a powerful and widely used approach for uncovering interpretable patterns in complex biological data. A well-known variant is non-negative matrix factorization (NMF) [11], which enforces non-negativity constraints on the factorized matrices. These constraints align naturally with biological data, such as gene expression levels, and enhance the interpretability of the learned components. NMF has been successfully extended to multi-omics integration, resulting in models such as IntNMF [12], PIntMF [5], GNMF [6], and iGMFNA [13]. These approaches aim to jointly analyze multiple omics layers and, in some cases, incorporate prior knowledge through biological interaction networks. However, most existing models rely on intra-omics relationships, such as within-layer features or sample similarities, while largely neglecting cross-omics feature interactions. This limitation hinders the ability to fully capture the regulatory complexity encoded across omics layers. To overcome this, a new matrix factorization framework is needed—one that integrates both intra- and cross-omics feature interactions while preserving interpretability and maintaining scalability for high-dimensional, multi-modal datasets.

In this work, we propose X-intNMF, a cross- and intra-omics network-regularized NMF model designed to integrate multi-omics data by incorporating both intra-omics and cross-omics feature interaction networks into a shared low-dimensional representation space. To enhance interpretability, the model imposes sparsity constraints on the omic-specific factor matrices. Unlike prior methods such as GNMF and iGMFNA, which primarily emphasize sample-sample interactions, X-intNMF leverages a feature-feature interaction network that captures both within-layer (intra-omics) and between-layer (cross-omics) relationships. This enables the model to incorporate known biological interactions, such as mRNA–miRNA regulatory networks, to improve integration quality. X-intNMF is an unsupervised learning framework applicable to diverse omics modalities, including mRNA, miRNA, and DNA methylation. It is also designed to be scalable and computationally efficient, making it suitable for large-scale multi-omics datasets.

## Materials and methods

This section introduces the X-intNMF framework, including the model formulation, optimization strategy, and the construction of cross- and intra-omics feature interaction networks. It also outlines the evaluation methods, baseline models, performance metrics, and parameter tuning process. All mathematical notations are summarized in Table S1 of the Supplementary document.

### Model description

Let **X**^(1)^, **X**^(2)^,…, **X**^(*D*)^ denote the input from *D* omics layers, where 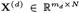, with *m*_*d*_ representing the number of features in the *d*^th^ omics layer and *N* denoting the number of samples. The proposed model is based on non-negative matrix factorization (NMF) [11], which aims to identify a shared non-negative sample factor matrix 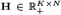 that captures sample-level factors, and *D* non-negative omic-specific factor matrices 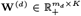 that project each omics layer into a *K*-dimensional latent space. Here, *K* is the pre-defined number of latent components. NMF seeks to optimize these matrices under the following objective:

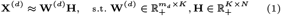

To ensure the interpretability of the model, sparsity constraints are imposed on both the sample factor matrix and the omic-specific factor matrices. These constraints promote more focused and biologically meaningful latent representations, resulting in the following formulation:

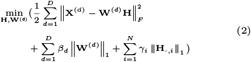

where *β*_*d*_ and *γ*_*i*_ are sparsity regularization parameters. The notation ‖*·* ‖_*F*_ denotes the Frobenius norm, while ‖*·* ‖_1_ refers to the *l*_1_ norm, and **H**_*·,i*_ represents the *i*^th^ column, corresponding the *i*^th^ sample, of the sample factor matrix **H**. However, standard NMF captures only internal structure within each omics layer and does not account for interactions within or between omics layers. To address this limitation, we propose an all-omics feature-feature interaction network **A**, defined as a *D × D* block matrix in Equation (3). The detailed procedure for constructing **A** matrix is described in Section *Intra- and cross-omics interaction network construction*.

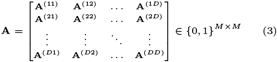

where **A**^(*pq*)^ denotes the adjacency matrix representing the interaction network between the *p*^th^ and *q*^th^ omics layers, and *M* = ∑*d m*_*d*_ represents the total number of features across all omics layers. The corresponding normalized adjacency matrix is given by **Ã** = **Δ**^*−*1*/*2^**AΔ**^*−*1*/*2^, where **Δ**^*−*1*/*2^ is the element-wise inverse square root of the degree matrix **Δ**, defined as:

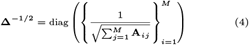

To incorporate this information into the model, a Laplacian regularization term 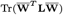 is introduced, where **L** is the normalized graph Laplacian matrix of the interaction network **A**, defined as:

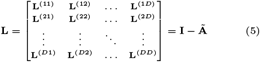

and 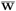 denotes the concatenation of all omic-specific factor matrices **W**^(*d*)^ along the feature axis (i.e., stacked vertically across *d*).

By the definition of the trace operator and the block structure of **L**, which follows the same *D × D* block partitioning as the interaction matrix **A**, the Laplacian regularization term can be reformulated as:

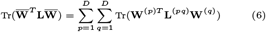

By partitioning the interaction network into blocks, interactions between omics layers can be computed directly from the data or derived from prior knowledge when available (e.g., mRNA–miRNA interaction networks). This design enhances the model’s robustness and adaptability across diverse omics datasets. By combining the NMF formulation with sparsity constraints (Equation (2)) and graph Laplacian regularization (Equation (6)), the final objective function of the model is defined as:

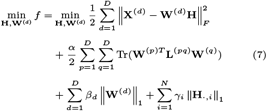

This objective function consists of four key components: the reconstruction error term, the cross- and intra-omics network regularization term, the omic-specific sparsity regularization term, and the sample-specific sparsity regularization term.

### Optimization strategy and convergence criteria

The objective function in Equation (7) is not jointly convex in all variables **W**^(*d*)^ and **H** and therefore must be optimized iteratively by updating one matrix while keeping the others fixed. The algorithm alternately updates the variables **W**^(*d*)^ and **H** until either a maximum number of iterations is reached or a convergence criterion is met. The convergence criterion is defined as the absolute change in the objective function falling below a pre-defined threshold. Further details on the iterative update formulas are provided in Section 2 of the Supplementary document.

For initialization, each **W**^(*d*)^ is first initialized using Non-negative Double Singular Value Decomposition (NNDSVD) as in standard NMF. The sample factor matrix **H** is then initialized based on the initialized **W**^(*d*)^ using the Lasso-based update rule from PIntMF [5]. When **W**^(*d*)^ is fixed, the objective function in Equation (7) reduces to Equation (8), which is solved with respect to each sample vector **H**_*·,i*_. This results in an independent Lasso problem for each *i* = 1,…,*N*.

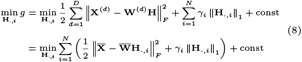

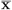 denotes the concatenation of all omics layers along the feature axis.

### Intra- and cross-omics interaction network construction

The interaction network **A** is structured as a block matrix, where each block **A**^(*pq*)^ is a binary submatrix representing interactions between omics layers *p* and *q*. This flexible design allows known interactions to be incorporated directly when available. In cases where computation is necessary, the interactions for a given omics pair (*p, q*)—whether intra- or cross-omics—are computed using the absolute value of the Pearson correlation coefficient (PCC) across all samples and then thresholded to match the average density of the existing interaction networks. This approach ensures that the resulting interaction network remains consistent with existing cross-omics interaction data, thereby preserving the biological relevance of the interactions.

### Evaluation methods

The performance of X-intNMF is evaluated based on its ability to classify disease outcomes, assess survival or disease-free time, and analyze cancer subtypes using the learned sample factor matrix **H**. The underlying assumption is that higher-quality representations derived from the multi-omics data in **H** will lead to improved predictive performance in downstream analyses.

#### Classification

In the binary classification tasks, the model is evaluated on two subtasks: (1) disease phenotype prediction and (2) classification of patients into pre-defined survival or disease-free duration categories (e.g., short-term vs. long-term survival). Performance is assessed using two commonly used metrics: Area Under the Receiver Operating Characteristic Curve (AUC) and Matthews Correlation Coefficient (MCC). AUC evaluates the model’s ability to distinguish between positive and negative classes across various threshold settings, making it robust to class imbalance. MCC considers all four confusion matrix categories—true positives, true negatives, false positives, and false negatives—and provides a more balanced and informative measure, particularly in imbalanced datasets.

#### Survival analysis

Beyond the binary classification of cancer phenotype and survival duration range, a separate survival analysis is conducted using a Cox proportional hazards model with an Elastic Net penalty [14]. This analysis aims to investigate the correlation between a patient’s overall survival, overall disease-free status, and the learned sample factor matrix **H**. The model optimizes the log-likelihood function *L*(***λ***) while incorporating both *l*_1_-norm and *l*_2_-norm penalties on the regression coefficients ***λ***, resulting in the following optimization problem:

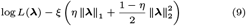

where *ξ ≥* 0 is the shrinkage control parameter, and *η ∈* [0, 1] is the mixing parameter that balances the *l*_1_-norm and *l*_2_-norm penalties. After training, the estimated coefficient ***λ*** is applied to the test set to compute the prognostic index (**PI**) for each patient:

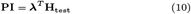

where **H**_test_ represents the dimensionality-reduced learned sample factor matrix of the test set. The median value of **PI** is then used as a threshold to classify patients into high-risk and low-risk groups. To evaluate the predictive performance, Kaplan-Meier survival curves are generated, and log-rank tests are performed to assess the statistical significance of survival or disease-free status differences between the high-risk and low-risk groups.

#### Cancer subtype analysis

Besides binary classification and survival analysis, the model is further evaluated based on its ability to cluster patients into different cancer subtypes. First, a sample-by-sample absolute Pearson correlation matrix is computed from the learned sample factor matrix **H**. Then, hierarchical clustering is performed to identify subtype-specific patterns. All clustering procedures are implemented using the scipy and seaborn libraries.

### Baselines

X-intNMF is compared with the following baselines, using their default configurations:

- **MOFA2** [4] is a Bayesian model for unsupervised multi-omics data integration. After dimensionality reduction, classification is performed using both Random Forest and Logistic Regression.
- **MOGONET** [9] is a supervised deep learning model for multi-omics data integration. It employs a View Correlation Discovery Network to capture cross-omics correlations in the label space. Unlike X-intNMF, MOGONET is supervised and does not incorporate known external cross-omics interaction networks.
- **MOMA** [15] is a supervised multi-task attention learning algorithm that uses a geometric approach to identify important modules in multi-omics data through an attention mechanism. The output probabilities of the two prediction tasks are averaged to generate the final classification.

- **MCRGCN** [8] is a supervised graph contrastive learning model for cancer subtype identification. It leverages the unique feature distributions of each omics layer and their interactions to enhance classification accuracy.

### Parameter tuning

The framework has four hyperparameters: the interaction network regularization parameter *α*, the number of latent components *K*, the omic-specific sparsity regularization parameters *β*_*d*_, and the sample-specific sparsity regularization parameters *γ*_*i*_. The parameters *γ*_*i*_ are computed directly during the initialization of **H** using Lasso cross-validation (LassoCV) from the *scikit-learn* package. The remaining parameters, *K, α*, and *β*_*d*_, are tuned via grid search in conjunction with a specific downstream task. In this study, hyperparameter optimization is performed based on the classification task. The resulting **H** from the best-performing configuration selected from the lists below is then reused for additional downstream analyses, including survival/disease-free outcome prediction and cancer subtype analysis.

- *K*: 10, 25, 50, 100, 200
- *α*: 0, 0.0001, 0.001, 0.01, 0.1, 1, 10, 100, 1000, 10000
- *β*_*d*_: 1, 0.1, 0.01, 10, 100 (applied uniformly for all *d*)
- Downstream classifiers: Logistic Regression (LR), Random Forest (RF)

For each classification target, 100 test cases are generated by randomly splitting the data into 80% training and 20% testing subsets. Within each test case, 5-fold cross-validation is performed on the training portion to evaluate all hyperparameter combinations. For each combination, the AUC scores across the five folds are averaged, and the set of parameters yielding the highest average AUC is selected. These optimal parameters are then used to train the model on the full training set of that test case, and evaluation, including AUC and MCC, is performed on the corresponding test set.

Regarding *ξ* and *η* in the Cox proportional hazards model, *η* is fixed at 0.5, while *ξ* is tuned using 5-fold cross-validation on the training set.

## Results

The performance of X-intNMF is evaluated through experiments on The Cancer Genome Atlas (TCGA) datasets, incorporating multi-omics profiles and interaction networks. This section begins with a detailed description of the datasets and data preprocessing procedures. Next, classification results are presented for both cancer phenotype prediction and survival duration classification, along with comparisons to baseline models. An ablation study is then conducted to assess the contribution of the interaction network to model performance. Additionally, results from survival and disease-free outcome analyses, as well as breast cancer subtyping, are reported to further demonstrate the effectiveness of the framework. Finally, the model’s scalability is evaluated through experiments in a three-omics setting.

### Data preparation

X-intNMF is evaluated on three TCGA datasets: breast cancer (BRCA), lung adenocarcinoma (LUAD), and ovarian cancer (OV). The omics layers used in the experiments include mRNA expression, miRNA expression, and DNA methylation, all obtained from the UCSC Xena Hub [16]. The mRNA-miRNA interaction network is sourced from TargetScanHuman [17]. The model is evaluated under two settings: (1) a two-omics setting using mRNA and miRNA expression, and (2) a three-omics setting that additionally includes DNA methylation.

All omics layers undergo preprocessing steps that include: removing samples not present across all omics layers, eliminating features with zero variance or all-zero values, and discarding mRNA/miRNA features not found in the interaction network. Due to the high dimensionality of the DNA methylation data, its features are further reduced by selecting the most variable features, limited to match the number of mRNA features after preprocessing. All retained samples are normalized by row-wise division using the maximum value in each row, scaling values to the [0, 1] range. The dimensions of the processed omics layers are summarized in Table 1.

**Table 1.** Number of features and samples after preprocessing for all datasets. The counts of mRNA, miRNA, and DNA methylation features are denoted as mRNA, miRNA, and DNAm, respectively.

|  | BRCA | LUAD | OV |
| --- | --- | --- | --- |
| **Feature#** |  |  |  |
| mRNA | 10,400 | 10,400 | 10,400 |
| miRNA | 277 | 277 | 274 |
| DNAm | 10,400 | 10,400 | 10,400 |
| **Sample#** |  |  |  |
| 2-omics | 830 | 466 | 304 |
| 3-omics | 674 | 456 | 295 |

For the breast cancer dataset, the classification targets include four phenotypes: Estrogen Receptor (ER+/ER-), Progesterone Receptor (PR+/PR-), Human Epidermal Growth Factor Receptor 2 (HER2+/HER2-), and Triple-Negative Breast Cancer (TN vs. non-TN). For the lung cancer and ovarian cancer datasets, the classification targets are survival and disease-free duration categories. Detailed information on the datasets and classification targets is provided in Table 2.

**Table 2.** Number of samples for each classification target. Counts are shown as “left/right,” indicating the number of samples in each of the two classes.

| Dataset | Target | 2-omics<br>sample# | 3-omics<br>sample# |
| --- | --- | --- | --- |
| BRCA | Phenotype: ER+/ER- | 185/54 | 153/45 |
|  | Phenotype: PR+/PR- | 159/80 | 130/68 |
|  | Phenotype: HER2+/HER2- | 39/200 | 28/170 |
|  | Phenotype: TN/non-TN | 46/193 | 40/158 |
| LUAD | Survival: $\leq 24 / \geq 30$ months | 93/134 | 93/134 |
| | Disease-free: $< 24 / \geq 24$ months | 54/99 | 54/99 |
| OV | Survival: $< 18 / \geq 18$ months | 40/207 | 38/202 |
| | Disease-free: $\leq 12 / \geq 18$ months | 17/71 | 15/68 |

### X-intNMF outperforms baselines in breast cancer phenotype classification

Table 3 summarizes the breast cancer phenotype classification results across competing methods. X-intNMF outperforms all baselines in both AUC and MCC across all four targets (ER, PR, HER2, and TN). In particular, X-intNMF achieves consistently high AUC values for ER, PR, and TN, demonstrating its robust ability to distinguish between positive and negative cases. Even for HER2, typically the most challenging target where many models underperform, X-intNMF achieves a strong AUC of 0.7656, outperforming all baseline methods. Although performance on HER2 is slightly lower compared to the other phenotypes, X-intNMF still provides reliable and accurate classification, underscoring its consistency across diverse prediction tasks.

**Table 3.** Breast cancer phenotype classification results. MOFA+RF and MOFA+LR represent the MOFA model combined with Random Forest and Logistic Regression classifiers, respectively. Bold values indicate the best performance for each metric.

| Methods | ER |  | PR |  | HER2 |  | TN |  |
| --- | --- | --- | --- | --- | --- | --- | --- | --- |
|  | MCC | AUC | MCC | AUC | MCC | AUC | MCC | AUC |
| X-intNMF | **0.7615** | **0.9612** | **0.7169** | **0.9232** | **0.2821** | **0.7656** | **0.6887** | **0.9652** |
| MOMA | 0.6990 | 0.9496 | 0.6613 | 0.9004 | 0.1022 | 0.7366 | 0.6252 | 0.9585 |
| MOGONET | 0.5730 | 0.9076 | 0.4372 | 0.8206 | 0.0134 | 0.6130 | 0.5266 | 0.9176 |
| MOFA+RF | 0.7500 | 0.9353 | 0.6798 | 0.8710 | 0.0759 | 0.5985 | 0.5816 | 0.9421 |
| MOFA+LR | 0.7544 | 0.9471 | 0.6688 | 0.8860 | 0.2318 | 0.6922 | 0.6430 | 0.9471 |
| MCRGCN | -0.0031 | 0.4961 | 0.0017 | 0.4954 | -0.0064 | 0.5051 | 0.0013 | 0.4897 |

In addition to its strong AUC performance, X-intNMF also achieves the highest MCC values among all methods, reflecting a better balance between sensitivity and specificity. For example, although MOMA attains competitive AUC scores for HER2, its MCC is notably lower than that of X-intNMF, further emphasizing the superior classification capability of the proposed framework. This improvement can be partly attributed to the effective integration of cross- and intra-omics interaction networks, which provide biologically meaningful context and enhance the quality of the learned representations.

### X-intNMF demonstrates robust performance in predicting survival-related duration range

In addition to cancer phenotype classification, X-intNMF is also evaluated on the task of predicting survival and disease-free duration categories for the lung and ovarian cancer datasets. Tables 4 and 5 present the results for the lung and ovarian cancer datasets, respectively.

**Table 4.**
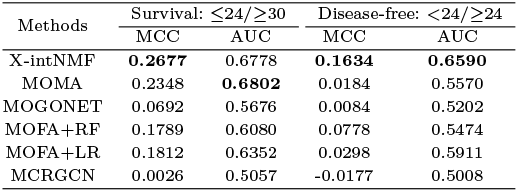
Results for lung cancer survival and cancer-free duration range classification. Bold values indicate the best performance for each task.

**Table 5.**
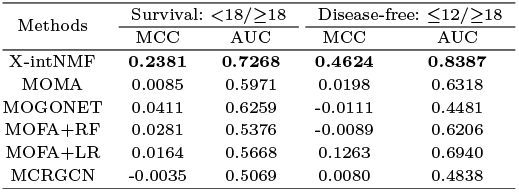
Results for ovarian cancer survival and cancer-free duration range classification. Bold values indicate the best performance for each task.

X-intNMF consistently outperforms baseline models in both survival and disease-free classification tasks for the two datasets, demonstrating strong performance in terms of both AUC and MCC. Although MOMA achieves the highest AUC for lung cancer survival prediction, X-intNMF surpasses it in MCC, indicating a more balanced and reliable performance. For the ovarian cancer dataset, X-intNMF outperforms all other models in both metrics, highlighting its robustness and accuracy in prognosis classification. These results suggest that X-intNMF is a powerful framework for predicting cancer outcomes, effectively leveraging multi-omics data to provide reliable and interpretable predictions.

### Impact of cross- and intra-omics feature interaction networks on model performance

To further validate the effectiveness of incorporating cross- and intra-omics feature interaction information into the learned multi-omics sample representations, an ablation study is conducted by setting the interaction network regularization parameter *α* to zero, which effectively disables the contribution of the interaction network during training. The results, presented in Table 6, illustrate the impact of this modification by comparing the classification performance of the ablated model with that of the full X-intNMF framework. Across all tasks, removing the interaction network leads to a notable decline in performance, particularly in terms of MCC. This decline is especially pronounced in complex and heterogeneous cases, such as HER2 classification in breast cancer and survival duration prediction in ovarian cancer. These findings confirm the importance of modeling cross- and intra-omics interactions in multi-omics data integration, demonstrating that incorporating such relationships enhances classification performance and robustness across diverse clinical contexts.

**Table 6.**
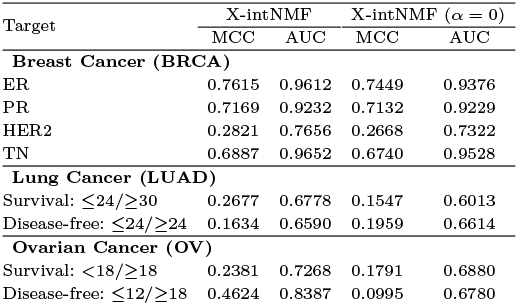
Classification results from the ablation study.

### X-intNMF demonstrates strong capability in survival analysis

In addition to classification tasks, survival analysis is conducted using the learned sample factor matrix **H** to evaluate overall survival and disease-free status, as described in the *Evaluation Methods* section. The analysis is performed on all three datasets and compared against both the ablated model (without the interaction network) and the concatenated original input data. Figures 2 and 3 show the Kaplan–Meier survival curves for the lung cancer dataset, illustrating differences in survival and disease-free status between high-risk and low-risk groups. X-intNMF achieves the most statistically significant log-rank test *p*-values and demonstrates the clearest separation between risk groups, outperforming both baselines in both analyses. These results highlight the robustness and clinical relevance of the learned representations. Additional results for the breast and ovarian cancer datasets are provided in the Supplementary document.

**Fig. 1.**
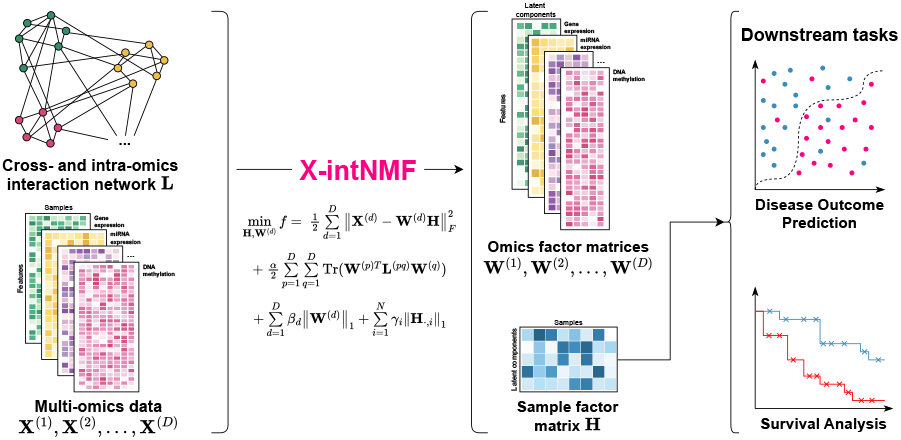
Overview of the proposed X-intNMF framework. X-intNMF integrates multi-omics data with both cross-omics and intra-omics interaction networks to jointly learn a shared sample factor matrix **H** and omic-specific factor matrices **W**^(*d*)^ via NMF with graph-based regularization. The resulting low-dimensional representations, **H** for samples and **W**^(*d*)^ for omics features, are then utilized for downstream analyses, including classification and survival prediction.

**Fig. 2.**
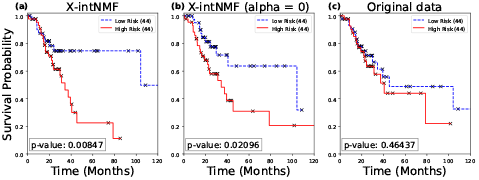
Kaplan–Meier survival analysis for overall survival in lung cancer. The input data for the Cox model in (a) is the sample factor matrix **H** learned by the proposed framework; (b) uses the sample factor matrix **H** from the ablated model with *α* = 0; and (c) uses the original input data.

**Fig. 3.**
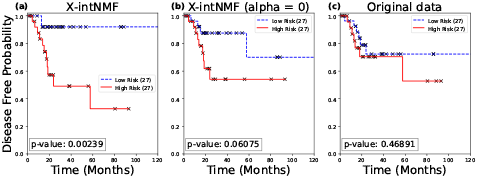
Kaplan–Meier survival analysis for disease-free status in lung cancer. The input data for the Cox model in (a) is the sample factor matrix **H** learned by the proposed framework; (b) uses the sample factor matrix **H** from the ablated model with *α* = 0; and (c) uses the original input data.

### X-intNMF effectively classifies breast cancer subtypes

In addition to the classification and survival analysis, the proposed framework is further evaluated for its ability to capture biologically meaningful subtypes using the breast cancer dataset. The subtype information, including Luminal A, Luminal B, HER2-enriched, and Basal-like, is obtained from the original TCGA study [18]. Figure 4 presents the clustermap of the absolute Pearson correlation matrix computed from the learned sample factor matrix **H**. The clustering patterns in the heatmap reveal that patient samples with the same subtype labels tend to group together, indicating strong within-subtype correlation. This trend is especially pronounced for the Basal-like subtype, which forms a distinct cluster. These results demonstrate that the learned low-dimensional representation from multi-omics data by X-intNMF can effectively capture subtype-specific signals.

**Fig. 4.**
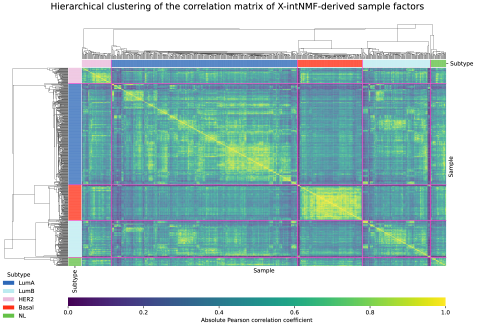
Clustermap of the absolute Pearson correlation matrix computed from the learned sample factor matrix **H**. The pink line marks the boundary between subtype labels. The subtype annotation bar indicates five subtypes: Luminal A (LumA), Luminal B (LumB), HER2-enriched (HER2), Basal-like (Basal), and Normal (NL).

### Scalability of X-intNMF beyond two omics modalities

To evaluate the framework’s compatibility and scalability, it is also tested using three omics layers, mRNA, miRNA, and DNA methylation, across all three datasets for both classification and survival analysis. Table 7 presents the classification results for survival and disease-free outcomes in the lung cancer dataset, using the same label definitions as in the two-omics setting (Table 2). The results demonstrate that the proposed framework continues to outperform baseline models in terms of both AUC and MCC, indicating that X-intNMF scales effectively to more complex multi-omics settings. Notably, MCRGCN yields near-random predictions on several targets in both the two- and three-omics settings, further underscoring the difficulty of the task and the relative strength of X-intNMF. Additional results and label definitions for the breast and ovarian cancer datasets are provided in the Supplementary document.

**Table 7.** Results for lung cancer survival and cancer-free duration range classification using the 3-omics setting. Bold values indicate the best performance for each task.

| Methods | Survival: $\leq 24 / \geq 30$ | | Disease-free: $\leq 24 / \geq 24$ | |
| --- | --- | --- | --- | --- |
|  | MCC | AUC | MCC | AUC |
| X-intNMF | **0.2758** | **0.6694** | **0.1934** | **0.6635** |
| X-intNMF$_{\alpha=0}$ | 0.2730 | 0.6568 | 0.0229 | 0.5204 |
| MOGONET | 0.1193 | 0.5820 | 0.0216 | 0.5456 |
| MOFA+RF | 0.1119 | 0.5717 | 0.0426 | 0.5588 |
| MOFA+LR | 0.0994 | 0.6102 | 0.0872 | 0.5972 |
| MCRGCN | -0.0217 | 0.5053 | 0.0091 | 0.4950 |

## Discussion

The results confirm that incorporating cross- and intra-omics feature interaction networks improves model performance across classification, survival analysis, and cancer subtyping tasks, underscoring the importance of capturing meaningful biological relationships. Moreover, the framework demonstrates strong compatibility with more than two omics layers, effectively scaling to three modalities and potentially beyond. However, hyperparameter selection remains a challenge. The current grid search approach requires extensive computational resources to identify optimal settings. Alternative optimization algorithms, such as particle swarm optimization [19], could reduce tuning time. Further improvements in data preprocessing and initialization strategies may also help stabilize learning dynamics and enhance model performance, robustness, and interpretability across diverse datasets.

While the framework achieves strong predictive performance, certain computational limitations persist. Objective function trajectories recorded via MLFlow (see Fig. S11 in the Supplementary document) show that the model converges rapidly during the first 1,000 iterations, but the convergence rate slows significantly thereafter. This suggests that the model may benefit from adaptive parameter schedules to accelerate convergence. Whether continued training beyond this point yields meaningful performance gains remains an open question. Although GPU acceleration is supported, optimization can still be slow, particularly for large datasets or when executed on CPUs. This bottleneck could potentially be addressed using methods such as the Alternating Direction Method of Multipliers (ADMM) [20], which may offer faster convergence and improved numerical stability.

The current construction of the cross- and intra-omics interaction network uses Pearson correlation with a fixed threshold for omics pairs lacking prior knowledge. However, more tailored strategies may be appropriate. Future work could explore adaptive interaction functions and thresholds specific to each omic-omic relationship, both within and across omics layers, to better capture the unique characteristics of each data modality. Similarly, the omic-specific sparsity regularization parameters *β*_*d*_ are currently set uniformly across layers; individualized tuning per omics type could further enhance performance. Another limitation is the exclusion of samples not present in all omics layers, which restricts the framework’s applicability. Future research should investigate methods for integrating such samples, including imputation techniques or approaches that explicitly handle missing data, enabling more inclusive and comprehensive analyses.

Lastly, while the sample factor matrix **H** is used for downstream tasks, the omic-specific factor matrices **W**^(*d*)^ remain underutilized, despite their potential value for feature selection, clustering, or pathway analysis. Future extensions of the framework could leverage **W**^(*d*)^ more fully to provide additional biological insights. Moreover, incorporating other data modalities, such as clinical records or imaging data, could broaden the framework’s utility, making it a more powerful and comprehensive tool for multi-omics integration.

## Conclusion

X-intNMF is a novel framework for multi-omics data analysis that integrates NMF with cross- and intra-omics feature interaction network regularization for cancer outcome classification, survival duration prediction, and cancer subtyping. The interaction network can be constructed using either prior biological knowledge or data-driven approaches, supporting both intra- and cross-omics relationships. This design enables the model to capture complex interdependencies across omics layers while maintaining the flexibility to adapt to diverse datasets. The framework is evaluated on three cancer types—breast, lung, and ovarian—demonstrating consistently improved performance over existing methods. Results further confirm that X-intNMF is compatible with and scalable to three omics layers, yielding meaningful insights in survival analysis and cancer subtype identification. Future work could explore leveraging the omic-specific factor matrices, investigate alternative initialization strategies, improve hyperparameter tuning through genetic algorithms, and evaluate the impact of different interaction functions and network density settings. Overall, X-intNMF provides a flexible, interpretable, and effective solution for integrative multi-omics analysis in cancer research.

## Notes

### Competing Interest Statement

The authors have declared no competing interest.

### Funding

- U.S. National Science Foundation: NSF-III2246796, NSF-III2152030

