## Supplementary for "X-intNMF: A Cross- and Intra-Omics Regularized NMF Framework for Multi-Omics Integration"

### Support Material

March 2025

#### 1 Notation Guideline

- All variables in **bold**, including those with a single index, represent either vectors or matrices. The bold numerals **0** and **1** denote zero and one vectors or matrices, depending on the context.
- All variables in non-bold form, including those with two indices (for matrices) or a single index (for vectors), represent scalar values.
- Superscripts enclosed in parentheses indicate indexing within a list or block of matrices. Specifically, a single integer in the superscript (e.g., $\mathbf{X}^{(d)}$) refers to the $d^{\text{th}}$ matrix in a list. A superscript with two integers (e.g., $\mathbf{X}^{(dp)}$) denotes the element at the $d^{\text{th}}$ row and $p^{\text{th}}$ column in a matrix block.

All mathematical notations used in the paper are summarized in Table S1.

| **Name** | **Type** | **Definition** |
|---|---|---|
| $D, N$ | Input | Number of omics layers, Number of samples |
| $m_d$ | Input | Feature size of $d^{\text{th}}$ omic layer |
| $M$ | Inferred | Total number of features |
| $\mathbf{X}^{(d)}$ | Input | $(N \times m_d)$-shaped matrix of $d^{\text{th}}$ omic layer |
| $\overline{\mathbf{X}}$ | Inferred | $(N \times M)$-shaped concatenation of all omics layers along the feature axis. |
| $\mathbf{A}$ | Input | $(M \times M)$-shaped adjacency matrix of the cross-intraomics feature interaction network |
| $\tilde{\mathbf{A}}$ | Inferred | $(M \times M)$-shaped normalized adjacency matrix of $\mathbf{A}$ |
| $\mathbf{\Delta}, \mathbf{\Delta}^{-1/2}$ | Inferred | $(M \times M)$-shaped degree matrix of $\mathbf{A}$ and its normalized form |
| $\mathbf{L}$ | Inferred | $(M \times M)$-shaped Laplacian matrix of the interaction network $\mathbf{A}$ |
| $\mathbf{A}^{(pq)}$ | Inferred | $(m_p \times m_q)$-shaped interaction network adjacency matrix between $p^{\text{th}}$ and $q^{\text{th}}$ omic layers |
| $\mathbf{L}^{(pq)}$ | Inferred | $(m_p \times m_q)$-shaped partition matrix of $\mathbf{L}$ |
| $\alpha$ | Hyperparameter | Interaction regularization parameter |
| $\beta_d$ | Hyperparameter | Omic-specific sparsity regularization parameters |
| $\gamma_i$ | Hyperparameter | Sample-specific sparsity regularization parameters |
| $K$ | Hyperparameter | Number of latent components |
| $\mathbf{W}^{(d)}$ | Output | $(m_d \times K)$-shaped $d^{\text{th}}$ omic factor matrix |
| $\mathbf{H}$ | Output | $(K \times N)$-shaped sample factor matrix |
| $\boldsymbol{\lambda}$ | Inferred | Regression coefficients in Cox proportional hazards model |
| $\xi, \eta$ | Hyperparameter | Shrinkage control and $l_1/l_2$-norm mixing parameters in Cox proportional hazards model |

**Table S1:** List of mathematical notations used in the paper.

### 2 Optimization of the X-IntNMF model

Since the objective function is not joinly convex over $\mathbf{W}^{(d)}$ and $\mathbf{H}$, it must be solved iteratively by optimizing one matrix while keeping the other fixed. Recall the the optimization problem of X-IntNMF:

$$\begin{aligned} \min_{\mathbf{H},\mathbf{W}^{(1)},\mathbf{W}^{(2)},...,\mathbf{W}^{(D)}} & f(\mathbf{H},\mathbf{W}^{(1)},\mathbf{W}^{(2)},...,\mathbf{W}^{(D)}) \\ \text{s.t. } & \mathbf{W}^{(d)} \in \mathbb{R}_+^{m_d \times K} \text{ for } d \in \{1,2,...,D\} \\ & \mathbf{H} \in \mathbb{R}_+^{K \times N}. \end{aligned} \tag{2.1}$$

where

$$\begin{aligned} f(\mathbf{H},\mathbf{W}^{(1)},\mathbf{W}^{(2)},...,\mathbf{W}^{(D)}) = & \frac{1}{2}\sum_{d=1}^{D}\left\|\mathbf{X}^{(d)} - \mathbf{W}^{(d)}\mathbf{H}\right\|_F^2 \\ & + \frac{\alpha}{2}\sum_{p=1}^{D}\sum_{q=1}^{D}\text{Tr}(\mathbf{W}^{(p)T}\mathbf{L}^{(pq)}\mathbf{W}^{(q)}) \\ & + \sum_{d=1}^{D}\beta_d\left\|\mathbf{W}^{(d)}\right\|_1 + \sum_{i=1}^{N}\gamma_i\left\|\mathbf{H}_{\cdot,i}\right\|_1 \end{aligned} \tag{2.2}$$

Let $\mathbf{\Phi}^{(d)} = [\phi_{ij}]^{(d)}$ and $\mathbf{\Psi} = [\psi_{ij}]$, the Lagrange multipliers $\mathcal{L}$ for $f$ is defined as (2.3):

$$\mathcal{L} = f + \text{Tr}(\mathbf{\Psi}\mathbf{H}^T) + \sum_{d=1}^{D}\text{Tr}(\mathbf{\Phi}^{(d)}\mathbf{W}^{(d)T}) \tag{2.3}$$

Equation (2.4) and (2.5) represent the partial derivatives of $\mathcal{L}$ with respect to $\mathbf{H} \in \mathbb{R}_+^{K\times N}$ and $\mathbf{W}^{(d)} \in \mathbb{R}_+^{m_d\times K}$:

$$\frac{\partial \mathcal{L}}{\partial \mathbf{W}^{(d)}} = (\mathbf{W}^{(d)}\mathbf{H} - \mathbf{X}^{(d)})\mathbf{H}^T + \alpha\sum_{p=1}^{D}\mathbf{L}^{(dp)}\mathbf{W}^{(p)} + \beta_d\mathbf{1} + \mathbf{\Phi}^{(d)} \tag{2.4}$$

$$\frac{\partial \mathcal{L}}{\partial \mathbf{H}} = \sum_{d=1}^{D}\mathbf{W}^{(d)T}(\mathbf{W}^{(d)}\mathbf{H} - \mathbf{X}^{(d)}) + \mathbf{\Psi} + \mathbf{1}\boldsymbol{\gamma} \tag{2.5}$$

Using Karush-Kuhn-Tucker (KKT) conditions $W_{ij}^{(d)}\phi_{ij}^{(d)} = 0$, $H_{ij}\psi_{ij} = 0$, $\frac{\partial \mathcal{L}}{\partial \mathbf{W}^{(d)}} = \mathbf{0}$, and $\frac{\partial \mathcal{L}}{\partial \mathbf{H}} = \mathbf{0}$, the update rules for $\mathbf{W}^{(d)}$ and $\mathbf{H}$ can be derived in Section 2.1 and 2.2 respectively.

#### 2.1 For omics factor matrices

From Equation (2.4) and condition of $\frac{\partial \mathcal{L}}{\partial \mathbf{W}^{(d)}} = \mathbf{0}$, the following equation can be derived:

$$(\mathbf{W}^{(d)}\mathbf{H} - \mathbf{X}^{(d)})\mathbf{H}^T + \alpha\sum_{p=1}^{D}\mathbf{L}^{(dp)}\mathbf{W}^{(p)} + \beta_d\mathbf{1} + \mathbf{\Phi}^{(d)} = \mathbf{0} \tag{2.6}$$

Swap $\mathbf{\Phi}^{(d)}$ to the right side and reverse the sign of Equation (2.6), the following equation can be derived:

$$\mathbf{\Phi}^{(d)} = (\mathbf{X}^{(d)} - \mathbf{W}^{(d)}\mathbf{H})\mathbf{H}^T - \alpha \sum_{p=1}^{D} \mathbf{L}^{(dp)}\mathbf{W}^{(p)} - \beta_d \mathbf{1} \tag{2.7}$$

Since $\mathbf{L}^{(dp)} = \mathbf{I}^{(dp)} - \tilde{\mathbf{A}}^{(dp)}$, Equation (2.7) can be rewritten as:

$$\begin{aligned}
\mathbf{\Phi}^{(d)} &= (\mathbf{X}^{(d)} - \mathbf{W}^{(d)}\mathbf{H})\mathbf{H}^T - \alpha \sum_{p=1}^{D} (\mathbf{I}^{(dp)} - \tilde{\mathbf{A}}^{(dp)})\mathbf{W}^{(p)} - \beta_d \mathbf{1} \\
&= (\mathbf{X}^{(d)} - \mathbf{W}^{(d)}\mathbf{H})\mathbf{H}^T + \alpha \sum_{p=1}^{D} (\tilde{\mathbf{A}}^{(dp)} - \mathbf{I}^{(dp)})\mathbf{W}^{(p)} - \beta_d \mathbf{1} \\
&= \left(\mathbf{X}^{(d)}\mathbf{H}^T + \alpha \sum_{p=1}^{D} \tilde{\mathbf{A}}^{(dp)}\mathbf{W}^{(p)}\right) - \left(\mathbf{W}^{(d)}\mathbf{H}\mathbf{H}^T + \alpha \sum_{p=1}^{D} \mathbf{I}^{(dp)}\mathbf{W}^{(p)} + \beta_d \mathbf{1}\right)
\end{aligned} \tag{2.8}$$

where $\mathbf{I}^{(dp)}$ is the block at row $d$ and column $p$ of the identity matrix. However, $\mathbf{I}^{(dp)}$ only non-zero when $d = p$, then:

$$\begin{aligned}
\mathbf{\Phi}^{(d)} &= \left(\mathbf{X}^{(d)}\mathbf{H}^T + \alpha \sum_{p=1}^{D} \tilde{\mathbf{A}}^{(dp)}\mathbf{W}^{(p)}\right) - \left(\mathbf{W}^{(d)}\mathbf{H}\mathbf{H}^T + \alpha \mathbf{W}^{(d)} + \beta_d \mathbf{1}\right) \\
&\equiv \mathbf{U}^{(d)} - \mathbf{V}^{(d)}
\end{aligned} \tag{2.9}$$

Combining Equation (2.9) with the KKT condition $W_{ij}^{(d)}\phi_{ij}^{(d)} = 0$, the following equation can be derived:

$$\Phi_{ij}^{(d)} W_{ij}^{(d)} = (U_{ij}^{(d)} - V_{ij}^{(d)}) W_{ij}^{(d)} = 0 \tag{2.10}$$

The update rule for $\mathbf{W}^{(d)}$ can be derived from Equation (2.10) as follows:

$$W_{ij}^{(d)} \leftarrow W_{ij}^{(d)} \times \frac{U_{ij}^{(d)}}{V_{ij}^{(d)}} = W_{ij}^{(d)} \times \frac{\left[\mathbf{X}^{(d)}\mathbf{H}^T + \alpha \sum_{p=1}^{D} \tilde{\mathbf{A}}^{(dp)}\mathbf{W}^{(p)}\right]_{ij}}{\left[\mathbf{W}^{(d)}\mathbf{H}\mathbf{H}^T + \alpha \mathbf{W}^{(d)}\right]_{ij} + \beta_d} \tag{2.11}$$

#### 2.2 For sample factor matrices

From Equation (2.5) and condition of $\frac{\partial \mathcal{L}}{\partial \mathbf{H}} = \mathbf{0}$, the following equation can be derived:

$$\sum_{d=1}^{D} \mathbf{W}^{(d)T}(\mathbf{W}^{(d)}\mathbf{H} - \mathbf{X}^{(d)}) + \mathbf{\Psi} + \mathbf{1}\boldsymbol{\gamma} = \mathbf{0} \tag{2.12}$$

Swap $\mathbf{\Psi}$ to the right side and reverse the sign of Equation (2.12), the following equation can be derived:

$$\begin{aligned}
\mathbf{\Psi} &= \sum_{d=1}^{D} \mathbf{W}^{(d)T}(\mathbf{X}^{(d)} - \mathbf{W}^{(d)}\mathbf{H}) - \mathbf{1}\boldsymbol{\gamma} \\
&= \left(\sum_{d=1}^{D} \mathbf{W}^{(d)T}\mathbf{X}^{(d)}\right) - \left(\sum_{d=1}^{D} \mathbf{W}^{(d)T}\mathbf{W}^{(d)}\mathbf{H} + \mathbf{1}\boldsymbol{\gamma}\right) \\
&\equiv \mathbf{S} - \mathbf{T}
\end{aligned} \tag{2.13}$$

Combining Equation (2.13) with the KKT condition $H_{ij}\psi_{ij} = 0$, the following equation can be derived:

$$\Psi_{ij} H_{ij} = (S_{ij} - T_{ij}) H_{ij} = 0 \tag{2.14}$$

The update rule for $\mathbf{H}$ can be derived from Equation (2.14) as follows:

$$H_{ij} \leftarrow H_{ij} \times \frac{S_{ij}}{T_{ij}} = H_{ij} \times \frac{\left[\sum_{d=1}^{D} \mathbf{W}^{(d)T}\mathbf{X}^{(d)}\right]_{ij}}{\left[\sum_{d=1}^{D} \mathbf{W}^{(d)T}\mathbf{W}^{(d)}\mathbf{H}\right]_{ij} + \gamma_i} \tag{2.15}$$

#### 2.3 Algorithm

The algorithm for the X-IntNMF model is summarized in Algorithm S1.

**Algorithm S1:** X-IntNMF algorithm

**Input:** $\mathbf{X}^{(d)}$, $\alpha$, $\beta_d$, $\gamma_i$, $K$

**Output:** $\mathbf{W}^{(d)}$, $\mathbf{H}$

Normalize $\mathbf{X}^{(d)}$ to $[0, 1]$ ;

Initialize $\mathbf{W}^{(d)}$ and $\mathbf{H}$;

**while** *not converged* **do**

  **for** $d = 1$ **to** $D$ **do**

    **for** $i = 1$ **to** $m_d$ **do**

      **for** $j = 1$ **to** $K$ **do**

        $W_{ij}^{(d)} \leftarrow W_{ij}^{(d)} \times \frac{\left[\mathbf{X}^{(d)}\mathbf{H}^T + \alpha \sum_{p=1}^{D} \tilde{\mathbf{A}}^{(dp)}\mathbf{W}^{(p)}\right]_{ij}}{\left[\mathbf{W}^{(d)}\mathbf{H}\mathbf{H}^T + \alpha\mathbf{W}^{(d)}\right]_{ij} + \beta_d}$;

      **end**

    **end**

  **end**

  **for** $i = 1$ **to** $N$ **do**

    **for** $j = 1$ **to** $K$ **do**

      $H_{ij} \leftarrow H_{ij} \times \frac{\left[\sum_{d=1}^{D} \mathbf{W}^{(d)T}\mathbf{X}^{(d)}\right]_{ij}}{\left[\sum_{d=1}^{D} \mathbf{W}^{(d)T}\mathbf{W}^{(d)}\mathbf{H}\right]_{ij} + \gamma_i}$;

    **end**

  **end**

**end**

**return** $\mathbf{W}^{(1)}, \mathbf{W}^{(2)}, ..., \mathbf{W}^{(D)}, \mathbf{H}$;

#### 3 3-omics classification experimental results

| Methods | ER | | PR | | HER2 | | TN | |
|---|---|---|---|---|---|---|---|---|
| | MCC | AUC | MCC | AUC | MCC | AUC | MCC | AUC |
| X-intMF | 0.6884 | **0.9515** | 0.6885 | **0.8973** | **0.2569** | **0.7472** | **0.6639** | **0.9594** |
| X-intNMF$_{\alpha=0}$ | 0.6848 | 0.9336 | 0.6659 | 0.8962 | 0.2075 | 0.6897 | 0.6569 | 0.9587 |
| MOGONET | 0.5300 | 0.8674 | 0.5445 | 0.8433 | -0.0049 | 0.5929 | 0.4173 | 0.8373 |
| MOFA+RF | 0.6990 | 0.9410 | 0.6682 | 0.8553 | 0.0230 | 0.6281 | 0.5429 | 0.9320 |
| MOFA+LR | **0.6995** | 0.9302 | **0.6908** | 0.8699 | 0.1355 | 0.7108 | 0.6215 | 0.9337 |
| MCRGCN | -0.0118 | 0.4861 | -0.0130 | 0.4858 | 0.0026 | 0.5390 | -0.0071 | 0.4980 |

**Table S2:** Result on 3-omics breast cancer phenotype classification. MOFA+RF and MOFA+LR are MOFA model combined with Random Forest and Logistic Regression classifiers. The best results are highlighted in bold.

| Methods | Survival: $<18/\geq 18$ | | Disease-free: $\leq 12/\geq 18$ | |
|---|---|---|---|---|
| | MCC | AUC | MCC | AUC |
| X-intMF | **0.1132** | **0.6505** | **0.3100** | **0.7210** |
| X-intNMF$_{\alpha=0}$ | -0.0266 | 0.5134 | 0.1703 | 0.6421 |
| MOGONET | 0.0137 | 0.5505 | -0.0130 | 0.4705 |
| MOFA+RF | -0.0064 | 0.5857 | -0.0121 | 0.5490 |
| MOFA+LR | -0.0040 | 0.6237 | 0.0888 | 0.6993 |
| MCRGCN | -0.0026 | 0.5070 | 0.0053 | 0.5105 |

**Table S3:** Result on ovarian cancer prognosis classification in terms of survival and cancer-free. MOFA+RF and MOFA+LR are MOFA model combined with Random Forest and Logistic Regression classifiers. The best results are highlighted in bold.

#### 4 Survival analysis result

The following figures present the Kaplan-Meier survival analysis results of the 2-omics and 3-omics datasets. All figures follow the same format: The input data used for the Cox model in (a) is the sample factor matrix $\mathbf{H}$ from the proposed framework, (b) is the sample factor matrix $\mathbf{H}$ from the ablated model with $\alpha = 0$, and (c) is from the original input data.

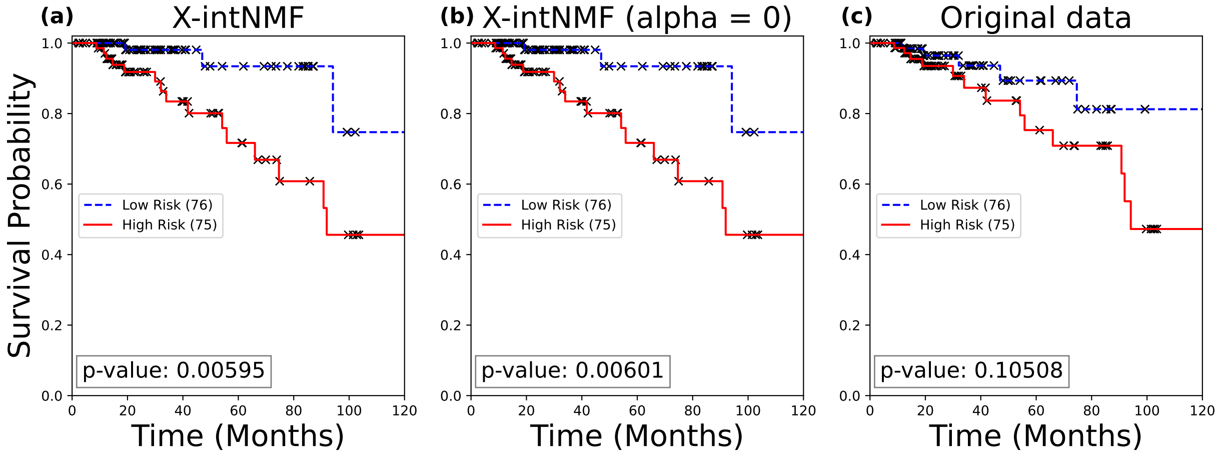


**Figure S1:** Kaplan-Meier survival analysis of the 2-omics breast cancer dataset.

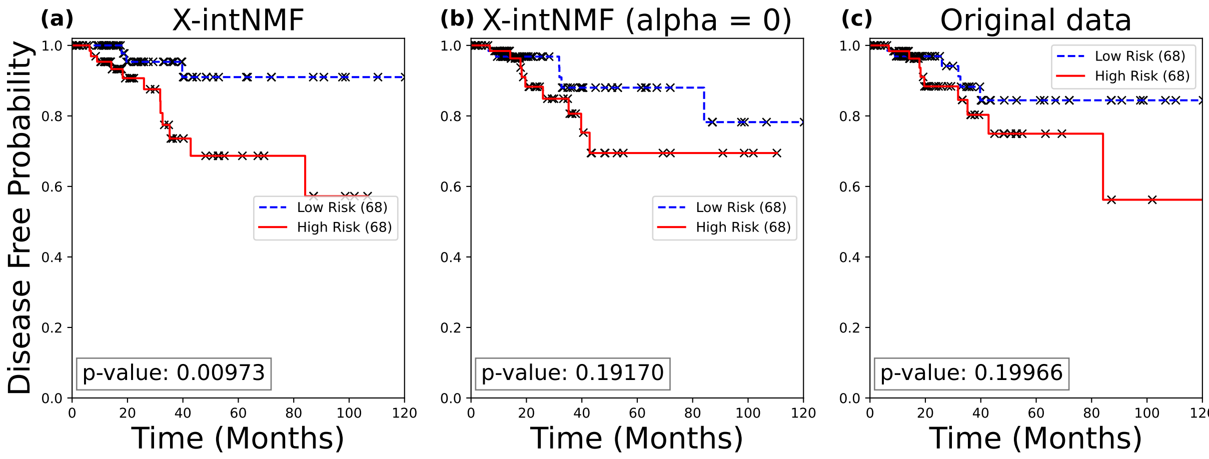


**Figure S2:** Kaplan-Meier disease-free analysis of the 2-omics breast cancer dataset.

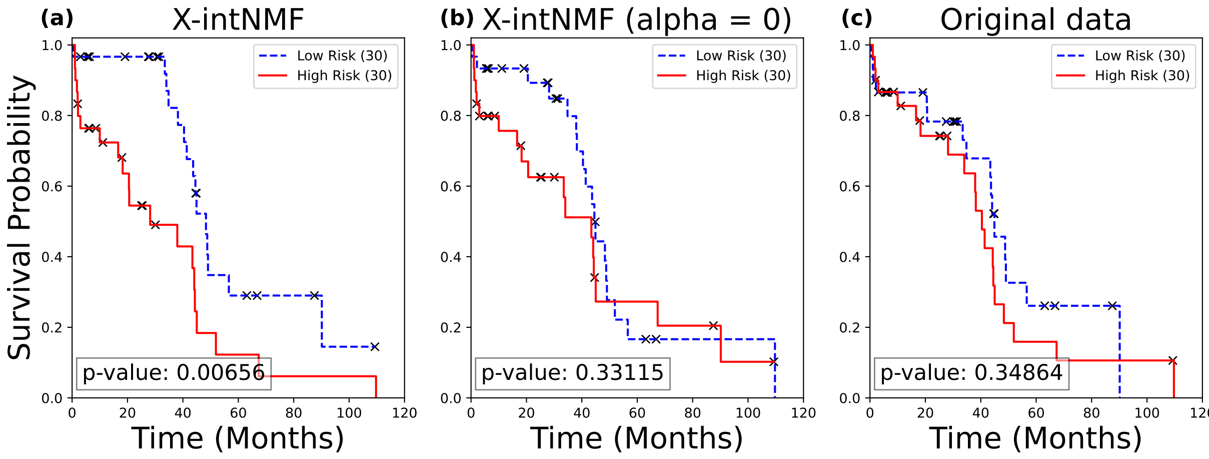


**Figure S3:** Kaplan-Meier survival analysis of the 2-omics ovarian cancer dataset.

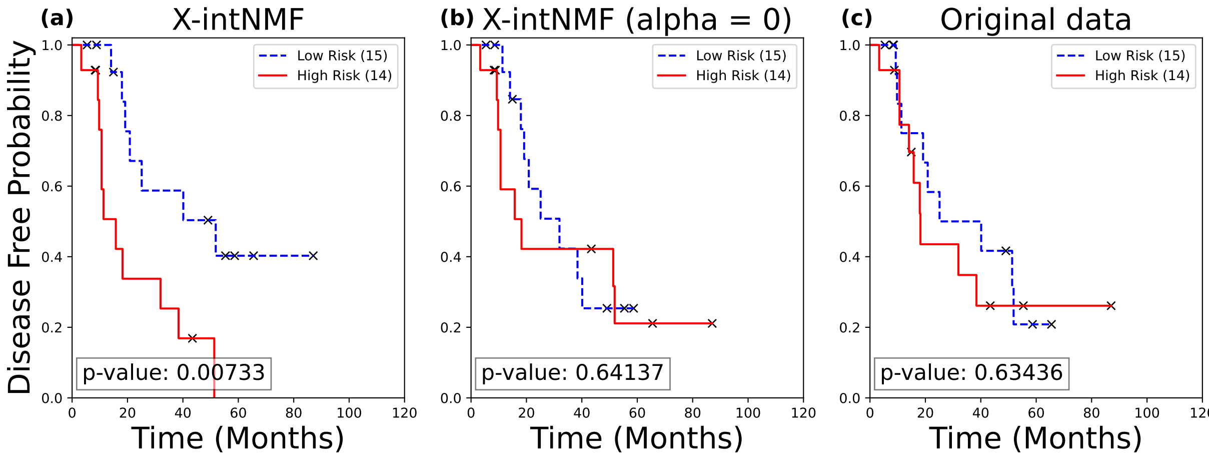


**Figure S4:** Kaplan-Meier disease-free analysis of the 2-omics ovarian cancer dataset.

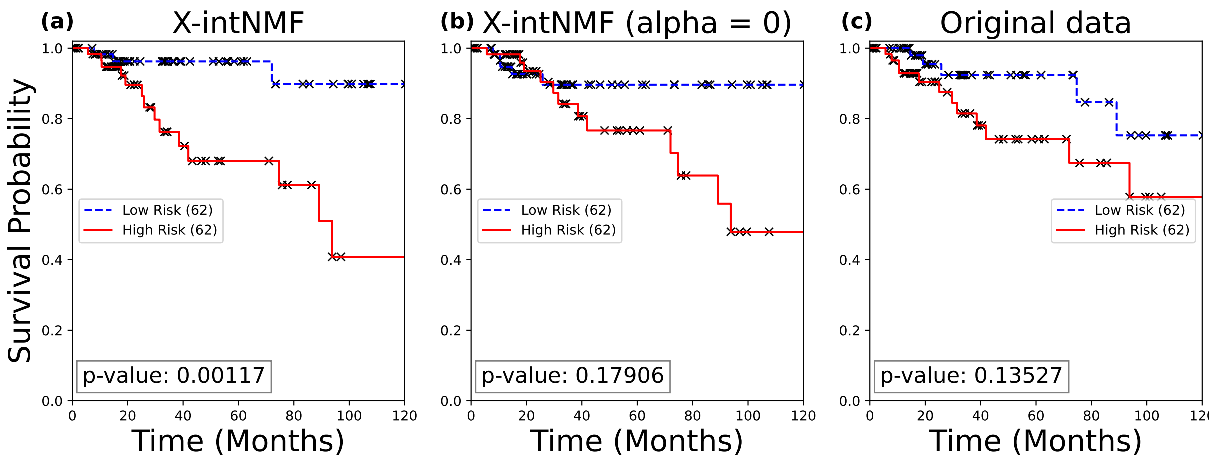


**Figure S5:** Kaplan-Meier survival analysis of the 3-omics breast cancer dataset.

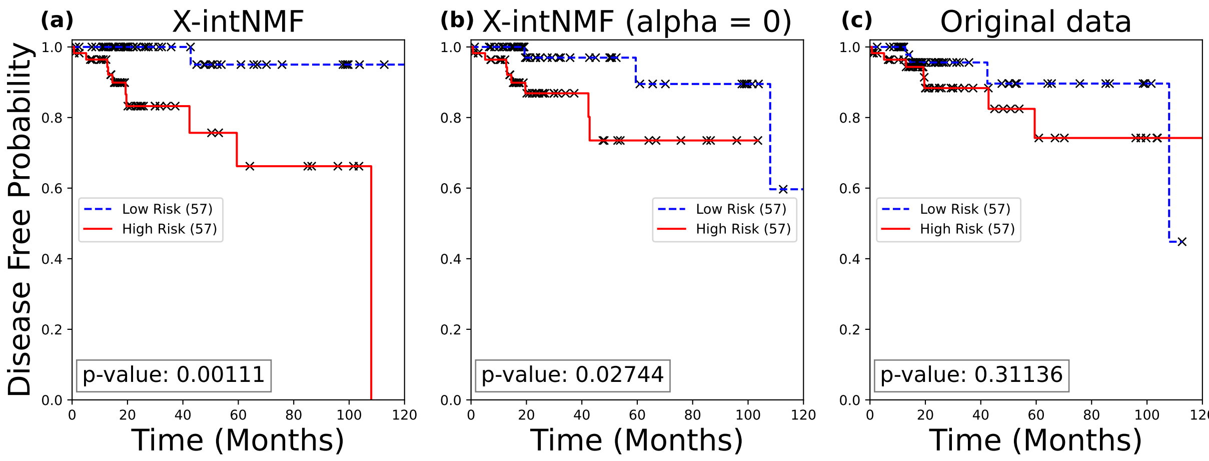


**Figure S6:** Kaplan-Meier disease-free analysis of the 3-omics breast cancer dataset.

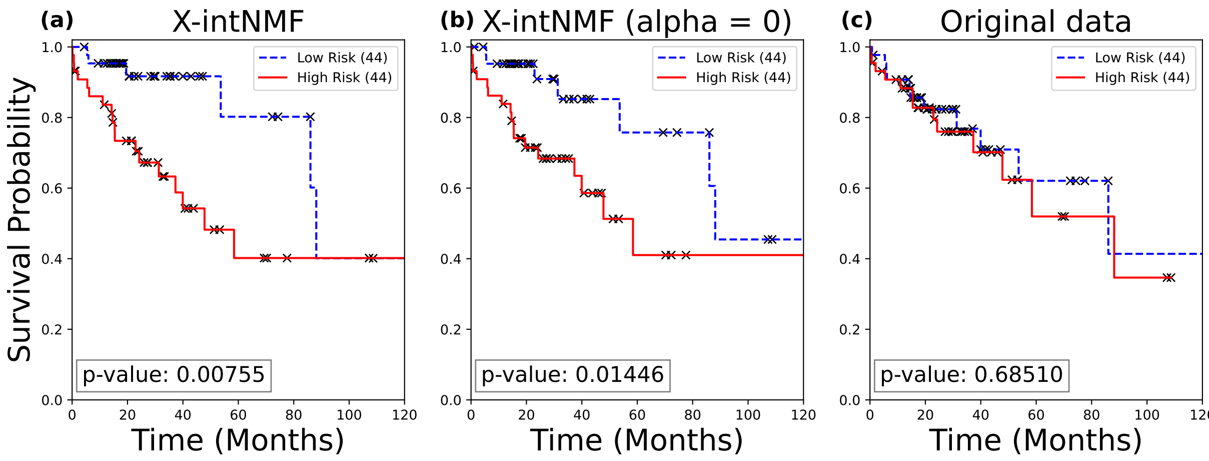


**Figure S7:** Kaplan-Meier survival analysis of the 3-omics lung cancer dataset.

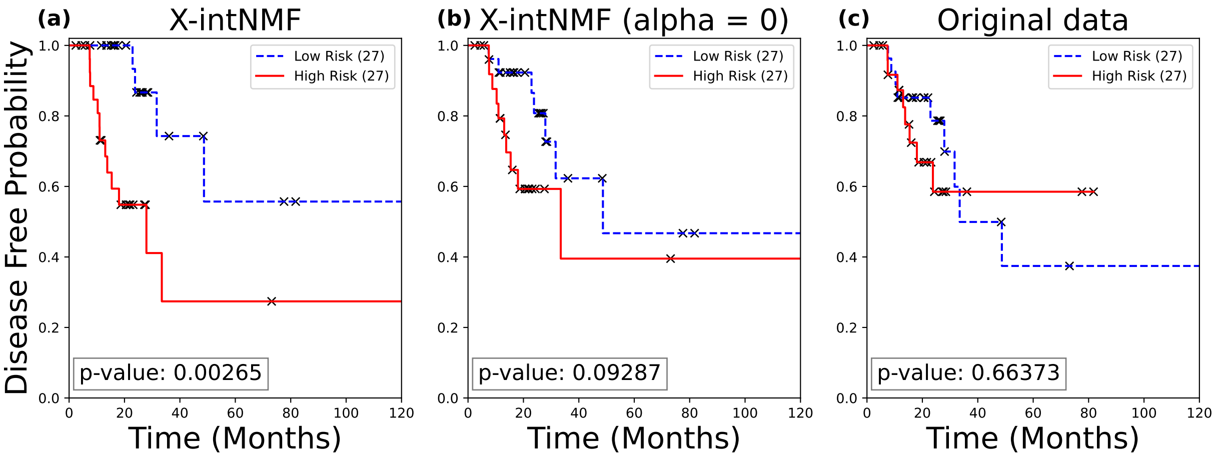


**Figure S8:** Kaplan-Meier disease-free analysis of the 3-omics lung cancer dataset.

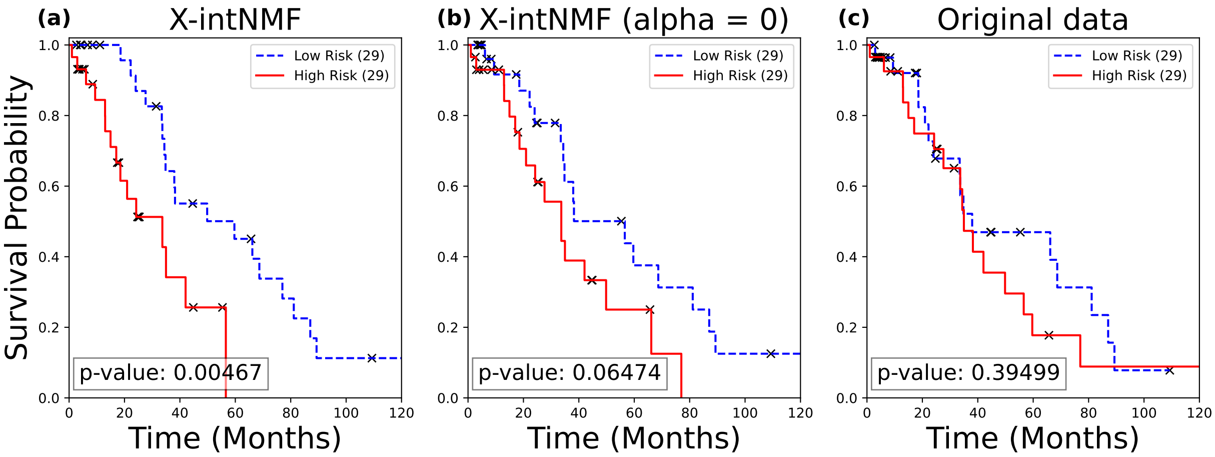


**Figure S9:** Kaplan-Meier survival analysis of the 3-omics ovarian cancer dataset.

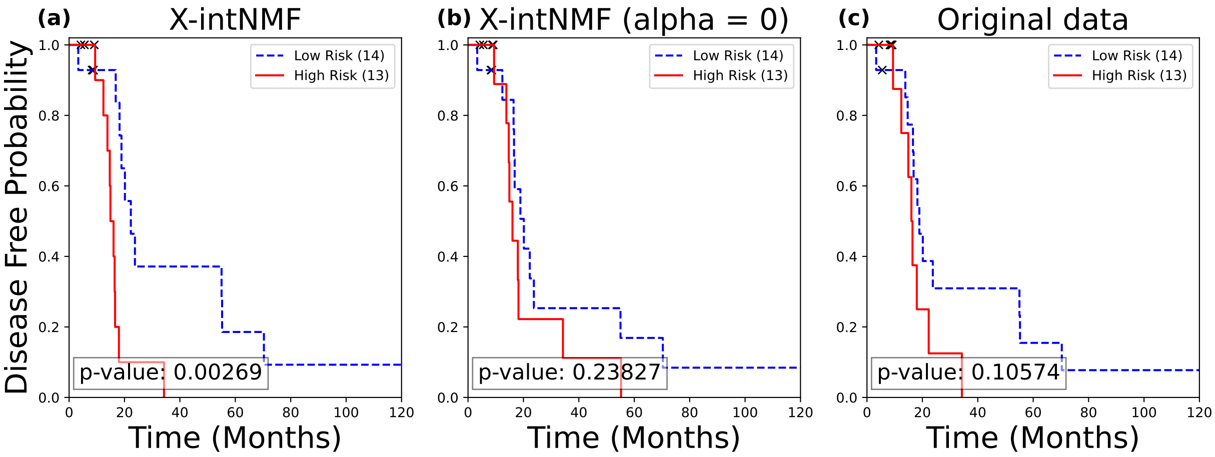


**Figure S10:** Kaplan-Meier disease-free analysis of the 3-omics ovarian cancer dataset.

### 5 Convergence logs over iterations

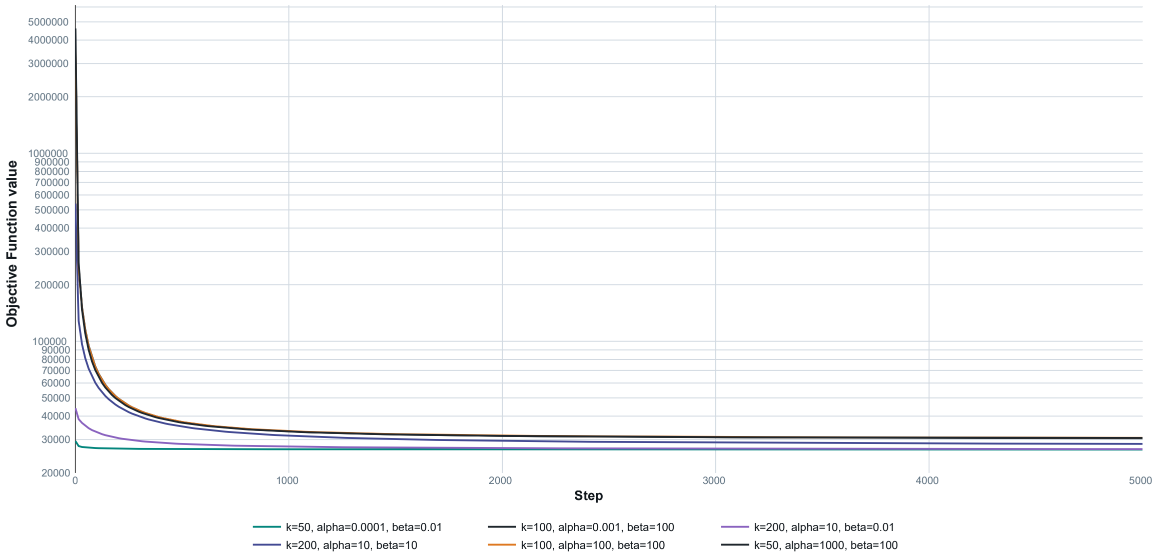


**Figure S11:** Convergence logs of the objective function under different parameter settings across iterations for the 2-omics breast cancer dataset, recorded using MLFlow.
